# PRSNet-2: End-to-end genotype-to-phenotype prediction via hierarchical graph neural networks

**DOI:** 10.1101/2025.11.22.689899

**Authors:** Han Li, Zisen Shan, Hongyu Fu, Yiwei Du, Shuaishuai Gao, Peng Gao, Johnathan Cooper-Knock, Yaosen Min, Xudong Xing, Sai Zhang

**Affiliations:** School of Mathematical Sciences and LPMC, Nankai University, Tianjin, China; Sheffield Institute for Translational Neuroscience, University of Sheffield, Sheffield, UK; China National Center for Bioinformation, Beijing, China; Beijing Institute of Genomics, Chinese Academy of Sciences, Beijing, China; College of Life Sciences, Nankai University, Tianjin, China; National Engineering Research Center of Ophthalmology and Optometry, Eye Hospital, Wenzhou Medical University, Wenzhou, China; Department of Environmental Health, Harvard T.H. Chan School of Public Health, Boston, USA; Zhongguancun Institute of Artificial Intelligence, Beijing, China; Department of Biomedical Informatics & Data Science, Yale University School of Medicine, New Haven, CT, USA

## Abstract

The emergence of large-scale biobanks has opened unprecedented opportunities for the development of data-driven approaches, especially deep learning-based methods, for genotype-to-phenotype (G2P) prediction. However, designing an end-to-end framework capable of directly leveraging extremely high-dimensional genotypic data while simultaneously mitigating overfitting and ensuring robust prediction remains a significant challenge. In this study, we introduce PRSNet-2, an interpretable end-to-end deep learning framework designed to predict complex phenotypes directly from large-scale genotypic data. PRSNet-2 proposes a novel hierarchical graph neural network (GNN) architecture, which first employs a multi-kernel aggregator to map high-dimensional genotypic features to gene-level representations. Next, it models gene-gene interactions through message-passing operations and uses an attention-based readout module to generate interpretable phenotypic predictions. We further introduce significance-guided regularization strategies to boost model’s generalizability based on prior genetic associations. Extensive empirical evaluations across multiple complex traits and diseases demonstrate that PRSNet-2 consistently outperforms a variety of baseline methods and exhibits superior capabilities in overcoming overfitting, even when modeling around half a million single nucleotide polymorphisms (SNPs). Moreover, PRSNet-2 is shown to be easily extendable to integrate multiple genome-wide association study (GWAS) datasets into a single model, thereby enhancing predictive performance for both single-phenotype and multi-phenotype prediction tasks. The inherent interpretability of PRSNet-2 further facilitates the identification of disease-relevant genes, functional gene modules, and potential therapeutic targets. In summary, PRSNet-2 offers a powerful, versatile tool for both genetic risk stratification and the discovery of biological insights.

## Introduction

The advent of large-scale biobanks, such as the UK Biobank (UKBB)^[1]^, FinnGen^[2]^, All of Us^[3]^, and the China Kadoorie Biobank^[4]^, which include extensive genetic data with detailed phenotypic information from hundreds of thousands to millions of individuals, has opened a new era for precision medicine research. These resources have enabled the application of data-driven approaches, especially deep learning methods that have demonstrated superior performance in big data modeling, to systematically uncover the genetic basis of complex traits and diseases, thereby achieving more accurate genotype-to-phenotype (G2P) prediction.

However, the exponential growth in both the volume and dimensionality of genetic data – often encompassing millions of SNPs per individual – poses significant challenges for deep learning-based G2P prediction. One of the biggest challenges is the curse of dimensionality, where the vast number of genetic variants, combined with the extreme sparsity of genetic signals (as most variants exert only small or negligible effects on phenotypes), can lead to overfitting and decreased model performance. Moreover, the limited availability disease labels – due to missing data or low prevalence of certain conditions – further complicates the modeling process and limits the model’s generalizability.

Traditional G2P prediction methods primarily rely on the polygenic risk score (PRS)^[5,6,7,8,9,10,11]^, which quantifies genetic susceptibility by aggregating the effects of genetic variants identified through the genome-wide association studies (GWAS)^[12,13,14]^. PRS-based methods, such as the clumping and thresholding (C+T) method^[5,15]^, PRSice-2 ^[6]^, LDpred2 ^[10]^, and lassosum2 ^[11]^, have shown promise in stratifying individuals at risk for diseases, guiding early intervention, and optimizing therapeutic strategies. However, these approaches typically follow an additive genetic architecture and oversimplify the complex relationships between genotype and phenotype. In an effort to capture more complex genetic interactions, we recently developed PRSNet^[16,17]^ (renamed from PRS-Net to simplify the nomenclature for subsequent versions), which calculates gene-level PRSs and uses graph neural networks (GNNs) to model non-additive genetic interactions. While PRSNet improves the predictive performance upon additive models, it requires time-consuming pre-computation of per-gene PRSs which reserves the additive structure at the gene level. Other methods, including tree-based models^[18,19,20,21,22,23]^ (e.g., random forests and gradient boosting) and deep learning models^[24,25]^ (e.g., multi-layer perceptrons), have also been proposed to account for non-linear genetic interactions. However, these models are often limited by the number of variants they can incorporate, leading to a significant loss of genetic information. Indeed, probably due to their inefficient modeling strategies, these non-linear approaches appear to be overfitting when dealing with the high-dimensional SNP data, exhibiting comparable or even worse performance compared to simpler additive methods^[16,17]^. Furthermore, many of these methods lack the integration of prior biological knowledge, which can diminish their prediction generalizability and interpretability. Therefore, there is an urgent need for an end-to-end deep learning framework that can effectively leverage high-dimensional genetic data from large biobanks while addressing overfitting and enhancing both predictive performance and biological interpretability.

In this study, we propose PRSNet-2, an interpretable end-to-end deep learning framework that predicts complex phenotypes directly from high-dimensional genotypic data. PRSNet-2 incorporates a novel hierarchical GNN architecture to layer-wise models SNP-to-gene, gene-to-gene, and gene-to-phenotype relationships. Specifically, PRSNet-2 first leverages a multi-kernel aggregator to map high-dimensional SNP features to gene-level representations, then models gene-gene interactions via message-passing operations, and finally employs an attention-based readout module to generate interpretable phenotype predictions. Moreover, we introduce significance-guided regularization strategies that integrate prior genetic associations to enhance model’s generalizability. Extensive empirical evaluations across eight complex diseases, including Alzheimer’s disease (AD), atrial fibrillation (AF), rheumatoid arthritis (RA), multiple sclerosis (MS), ulcerative colitis (UC), abdominal aortic aneurysm (AAA), coronary artery disease (CAD) and asthma, demonstrate that PRSNet-2 consistently outperforms a wide range of baseline methods, including both traditional PRS approaches and machine learning models. Evaluations on an independent test cohort further highlight the superior generalizability of PRSNet-2 across diverse cohorts. PRSNet-2 also excels in predicting complex traits such as height and body mass index (BMI) compared to baseline methods. Furthermore, PRSNet-2 is highly extendable, allowing for the integration of multiple GWAS datasets, thereby enhancing prediction performance for both single-phenotype and multi-phenotype prediction tasks via a multitask learning framework. Across all prediction tasks, PRSNet-2 exhibits robust performance, effectively mitigating overfitting when modeling high-dimensional SNP datasets containing up to half a million variants. The inherent interpretability of PRSNet-2 further facilitates the identification of disease-relevant genes, functional gene modules, and potential therapeutic targets. Collectively, PRSNet-2 provides a powerful and versatile tool for concurrent genetic risk stratification and biological insight discovery.

## Methods

In this section, we present our PRSNet-2 for genotype-to-phenotype prediction, which includes the construction of the hierarchical graph, the detailed architecture of the hierarchical GNN, the significance-guided regularization strategies, and its scalability to integrate multiple GWAS datasets (Fig. 1).

**Fig. 1:**
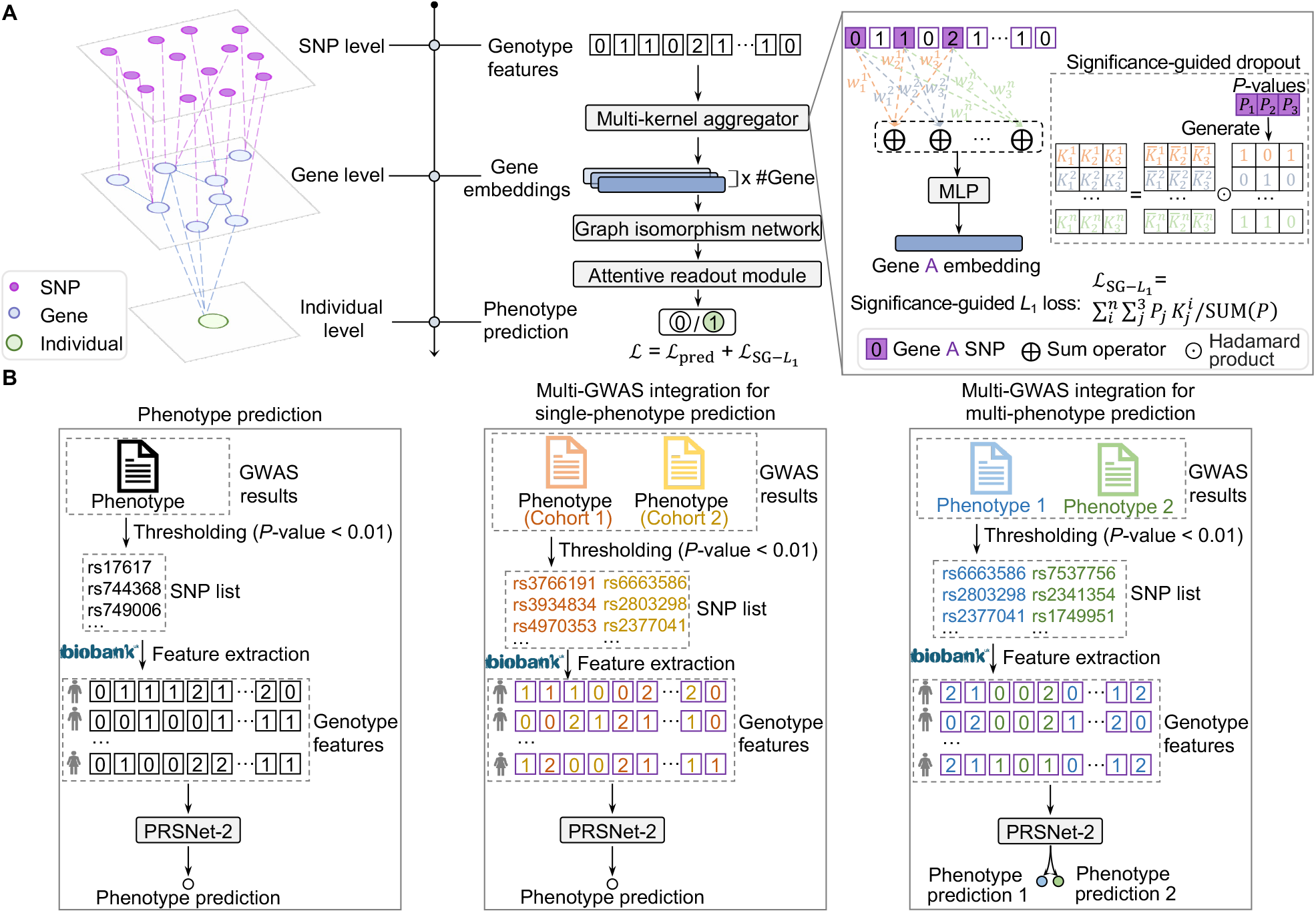
An illustrative diagram of PRSNet-2. (A) PRSNet-2 is an end-to-end genotype-to-phenotype prediction framework based on a graph neural network that hierarchically models SNP-to-gene, gene-to-gene, and gene-to-phenotype relationships. PRSNet-2 first employs a multi-kernel aggregator to map high-dimensional genotypes to gene-level representations, then captures gene-gene interactions through a graph isomorphism network, and finally uses an attention-based readout module to generate interpretable phenotypic predictions. Regularization strategies are introduced to mitigate overfitting, including significance-guided dropout in the multi-kernel aggregator and significance-guided *L*_1_ penalty term in the loss function. (B) PRSNet-2 can be applied in multiple prediction scenarios, including: (1) single-phenotype prediction using a single GWAS, where summary statistics from one GWAS are used to select a set of significant SNPs and then generate genotype features (coded as 0, 1, and 2, representing allele dosages); (2) single-phenotype prediction using multiple GWASs, where summary statistics from several GWASs are combined to select a list of significant SNPs; and (3) multi-phenotype prediction, where summary statistics from multiple corresponding GWASs are integrated to select significant SNPs for phenotype prediction utilizing a multitask learning strategy. SNP, single-nucleotide polymorphism; GWAS, genome-wide association study; MLP, multi-layer perceptron.

### Hierarchical graph construction

To model the hierarchical genetic structure underlying a specific phenotype, we first construct a hierarchical, heterogeneous graph for each individual (Fig. 1A), denoted as 𝒢 = (𝒱, E), where 𝒱 represents the set of nodes and ℰ represents the edges between nodes. The graph consists of three types of nodes: SNP nodes (denoted as 𝒱_SNP_), gene nodes (denoted as 𝒱_G_), and the phenotype node.

The SNP nodes are derived from GWAS summary statistics, where SNPs are selected on the basis of a *P* -value threshold, with the default set to 0.01. Only SNPs with a *P* -value below this threshold are retained. The feature value of each SNP node is encoded as 0, 1, or 2, corresponding to the allele dosage (i.e., 0/0, 0/1, and 1/1).

Gene nodes represent the 18,963 protein-coding genes in the human genome. An edge is established between a SNP and a gene if the SNP falls within the gene body extended by 10 kb both upstream and downstream. This allows the graph to capture the relationships between genetic variants and the genes they potentially affect. Additionally, we introduce a gene-gene interaction (GGI) network, where edges are formed between gene nodes based on high-confidence protein-protein interactions (PPIs) derived from the STRING database^[26]^. Only interactions with a confidence score greater than 0.8 are considered. This step enhances the gene-level graph by incorporating functional relationships between genes.

Finally, a phenotype node is connected to all gene nodes in the graph, which is used to summarize the genetic information from the gene nodes for phenotype prediction.

Note that PRSNet-2 uses two cohorts: the base cohort, which is analyzed in GWAS to estimate per-variant effect sizes and *P* -values, and the target cohort, which is used to train and test PRSNet-2. To prevent information leakage, we selected GWAS data that is independent of the UKBB for each disease and trait.

### PRSNet-2

The PRSNet-2 framework includes a novel hierarchical GNN architecture to model the SNP-gene-phenotype graph. The GNN architecture consists of three key components: the multi-kernel aggregator, the graph isomorphism network, and the attentive readout module (Fig. 1A).

#### Multi-kernel aggregator

The multi-kernel aggregator is designed to encapsulate high-dimensional genotypic features by mapping SNP nodes to gene-level representations through an ensemble of multiple kernels. Specifically, for each gene node 𝒱_*g*_ ∈ 𝒱_G_ and its associated SNPs 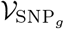 (i.e., SNPs linked to the gene *g*), the gene-level representation (denoted as ***h***_*g*_) is computed as follows:

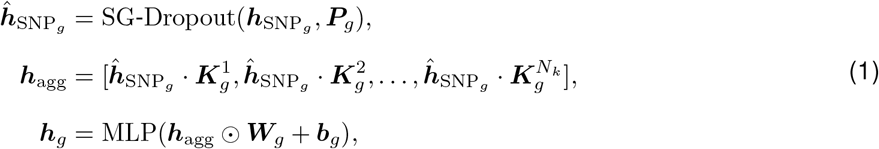

where 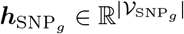 represents the genotype features for all SNP nodes linked to gene *g*, 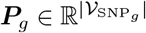 denotes the *P* -values derived from the GWAS summary statistics for corresponding SNPs, and 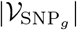 stands for the total number of SNPs associated with gene *g*. Each element of 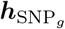 corresponds to the allele dosage of the respective SNP. The function SG-Dropout(·) represents a significance-guided dropout mechanism that modifies the feature vector 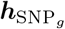 by selectively dropping out SNPs based on their statistical significance, as derived from GWAS. Specifically, the dropout probability for SNP *i* (denoted as *p*_*i*_) is computed as follows:

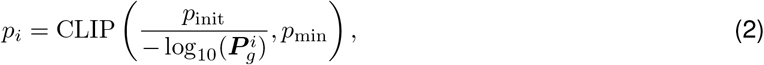

where CLIP(·) denotes a clipping function, 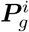 is the GWAS *P* -value for SNP *i, p*_init_ is an initial scaling factor for the dropout rate, and *p*_min_ is the minimum allowable dropout probability. The dropout probability for each SNP is inversely related to its GWAS *P* -value, ensuring that more significant SNPs (with lower *P* -values) are less likely to be dropped out during training. This dynamic dropout strategy regularizes the learning process and enhances the model’s capacity to focus on more relevant genetic variants.

Each kernel 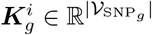 represents a learnable transformation for gene *g*, and *N*_*k*_ denotes the number of kernels. Using an ensemble of multiple kernels, the aggregator aggregates diverse transformations of the SNP features, allowing for a richer and more comprehensive characterization of the genetic information.

After aggregating the transformed SNP features across all kernels, the resulting feature vector is passed through a learnable scaling transformation, where 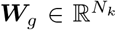 and 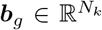 are the learnable weight and bias vectors for gene *g*, respectively. The operator ⊙ denotes the Hadamard product (i.e., element-wise multiplication). The scaled feature vector is then processed through a multi-layer perceptron (MLP), which maps the feature dimension from *N*_*k*_ to *D*. The output, ***h***_*g*_ ∈ ℝ^*D*^, is the derived feature vector for gene *g*.

This process is repeated for all genes, yielding the overall feature representation matrix 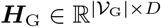, where *D* is the dimension of the latent feature space and |𝒱_G_| is the total number of genes.

In summary, the multi-kernel aggregator integrates SNP-level features into gene-level representations through an ensemble of learnable kernels and significance-guided dropout. This approach is designed to enhance the model’s ability to capture complex genotypic patterns and interdependence across genes, while mitigating the influence of noisy genetic factors.

#### Graph isomorphism network

We utilize a graph isomorphism network (GIN)^[27]^ to model GGIs due to its demonstrated theoretical and empirical expressiveness. In particular, given gene node features ***h***_*g*_ ∈ ***H***_G_ for *v*_*g*_ ∈ 𝒱_G_, we apply multiple GIN layers to iteratively update the features of each node by aggregating neighboring information, as depicted below:

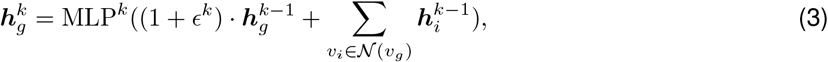

where 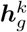 is the hidden feature of *v*_*g*_ at the *k*-th layer, 𝒩 (*v*_*i*_) stands for the neighbors of *v*_*i*_ in the GGI network, MLP^*k*^ is the MLP at the *k*-th layer, and *ϵ* is a learnable variable. After *N*_*L*_ steps of message passing, each gene encapsulates the genetic information within its *N*_L_-hop neighborhoods, and the derived gene embeddings are denoted as 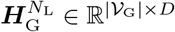.

#### Attentive readout module

Following the GNN operation, we compute a global representation for each individual (denoted as ***h***_𝒢_) using an attentive readout module shown as follows:

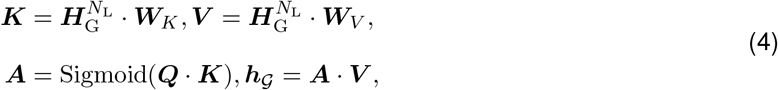

where ***W***_*K*_ ∈ ℝ^*D*×*D*^ and ***W***_*V*_ ∈ ℝ^*D*×*D*^ are trainable key (i.e., ***K***) and value (i.e., 𝒱_G_) matrices, respectively, ***Q*** ∈ ℝ^*D*^ is the trainable query vector, ***A*** ∈ ℝ^|**𝒱**|^ are the attention scores, and ***h***_𝒢_ ∈ ℝ^*D*^ is the global representation. Note that a higher attention score indicates a greater disease-relevance of the corresponding gene.

Using the global representation ***h***_𝒢_, we then apply an MLP as a predictor to generate the final prediction, denoted as *ŷ*. For binary phenotype (e.g., disease) prediction, we employ a cross-entropy loss (*L*_pred_), whereas for quantitative trait prediction, we use a mean squared error (MSE) loss.

To further improve model generalization, we introduce a significance-guided *L*_1_ loss (denoted as *L*_SG−L1_), defined as:

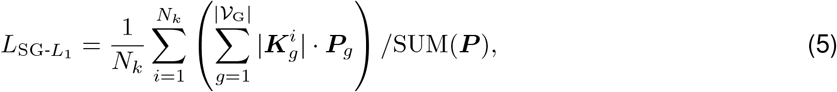

where 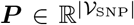 is the *P* -value vector for all SNPs, and ***P***_*g*_ is the *P* -value vector for gene *g*. This loss function encourages sparsity in the learned kernel values while taking into account the GWAS significance of SNPs. The aforementioned dropout strategy is designed to reduce the impact of less significant variants, while this *L*_1_ loss forces the model to focus on the most associated SNPs, together preventing overfitting to noise or irrelevant variants.

The final loss is defined as the sum of the predictive loss and the *L*_1_ regularization:

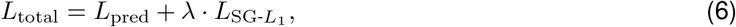

where *λ* is the regularization weight.

#### Multi-GWAS integration

PRSNet-2 can be extended to integrate multiple GWASs, leveraging diverse genetic signals across studies to improve both predictive accuracy and robustness (Fig. 1B). In the case of integrating two GWASs, the hierarchical graph 𝒢 will contain two sets of SNPs: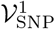 and 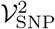, which represent the SNPs selected based on the first and second GWAS respectively. The multi-kernel aggregator is modified to handle these two sets of SNPs as follows:

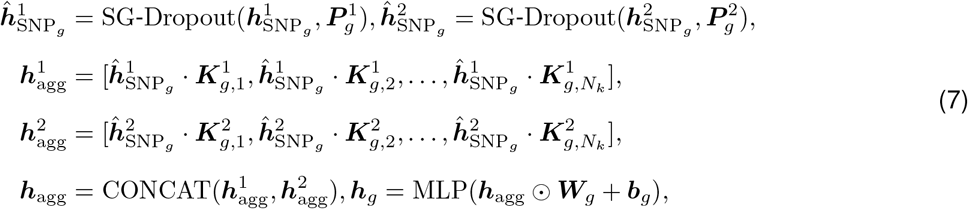

where 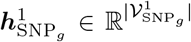 and 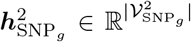 represent the genotype feature vectors for all SNP nodes linked to gene *g* selected from the first and second GWAS, respectively. 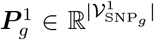 and 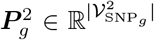 are the GWAS *P* -values associated with the SNPs from the first and second GWAS results, respectively. 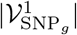 and 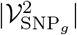 denote the numbers of SNPs. 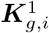 and 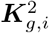 are the learnable kernels for gene *g* corresponding to the first and second GWAS, respectively.

After obtaining the aggregated SNP features for both GWASs, we concatenate the features and pass them through an MLP to generate the gene-level feature vector ***h***_*g*_ for gene *g*.

#### Single phenotype prediction

If we have two GWASs for the same phenotype, the significance-guided *L*_1_ loss function is adapted to account for both GWASs:

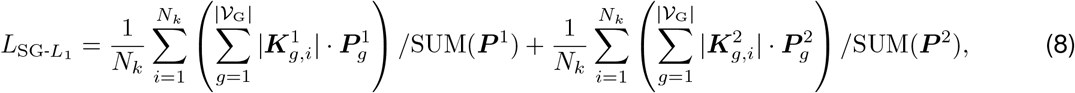

where ***P*** ^1^ and ***P*** ^2^ are the *P* -value vectors for all SNPs selected from the first and second GWAS, respectively. SUM(***P*** ^1^) and SUM(***P*** ^2^) denote the normalization terms.

#### Multiple phenotype prediction

Assume we have two GWASs for two different phenotypes, we use a multitask learning framework to simultaneously model both phenotypes. In this case, we pass the integrated gene representations 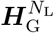 into two independent attentive readout modules and MLP layers for phenotype prediction. The total loss for this scenario is the sum of the predictive losses for both phenotypes and the significance-guided *L*_1_ loss:

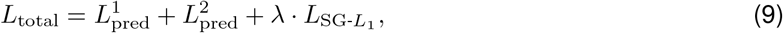

where 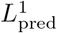 and 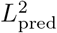 are the predictive losses for the first and second phenotypes, respectively.

Note that PRSNet-2 can be defined in the same way in the case of more than two GWASs.

## Results

### PRSNet-2 improves disease risk prediction

We first evaluated the performance of PRSNet-2 in predicting disease risk. Genotype and phenotype data were extracted from the UK Biobank database for eight complex diseases: Alzheimer’s disease, atrial fibrillation, rheumatoid arthritis, multiple sclerosis, ulcerative colitis, abdominal aortic aneurysm, and asthma. Disease endpoints were defined using the ICD-10 codes (Supplementary Table 1). For the primary analysis, we focused exclusively on individuals of Western European ancestry, as the non-European population was insufficiently represented to provide adequate training and test data (Supplementary Table 2). After quality control, each disease dataset comprised approximately 411,000 individuals (Supplementary Methods). These datasets were randomly partitioned into training, validation, and test sets with an 8:1:1 ratio. To benchmark PRSNet-2’s performance, we compared it with several baseline methods: traditional PRS methods including PLINK, PRSice2, lassosum2, and LDpred2; machine learning-based methods including MLP and XGBoost; and its predecessor PRSNet. We assessed model’s performance using two metrics: the area under the receiver operating characteristic curve (AUROC) and the area under the precision-recall curve (AUPRC). To ensure robustness and reliability, we performed five independent runs with different random seeds for each model and each disease.

PRSNet-2 consistently outperformed all baseline methods across all eight diseases in both AUPRC and AUROC (Fig. 2A). At the individual disease level, PRSNet-2 demonstrated superior performance, surpassing all baseline methods in AUPRC for all diseases and in AUROC for seven diseases (Fig. 2A). Given the high imbalance of these disease datasets, PRSNet-2 achieved greater relative improvements in AUPRC compared to AUROC (Fig. 2B and Supplementary Fig. 1). Notably, PRSNet-2 showed the greatest relative improvement of 4.76% for multiple sclerosis and 4.42% for Alzheimer’s disease, in comparison with the best baseline method (i.e., PRSNet). We also evaluated the generalizability of PRSNet-2 across different cohorts using the AD Neuroimaging Initiative (ADNI)^[28]^. PRSNet-2 was trained on the original target cohort (i.e., UKBB), and all PRS methods were tested on the independent ADNI cohort. PRSNet-2 continued to outperform all baseline methods in terms of both AUROC and AUPRC (Fig. 2C). These results underscore the superior performance and generalizability of PRSNet-2 in disease risk prediction.

**Fig. 2:**
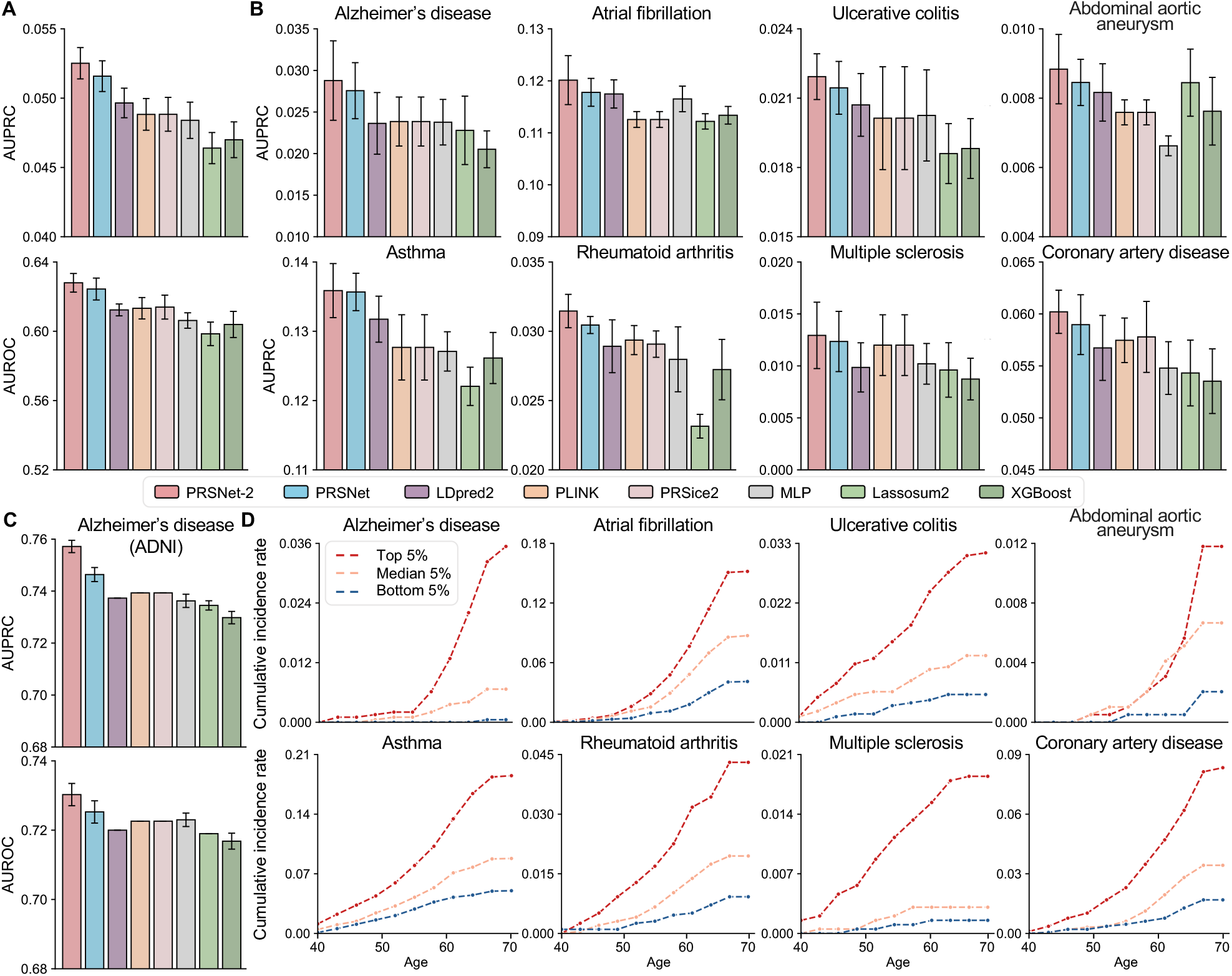
Performance evaluation for disease prediction. (A) The average AUPRC and AUROC scores of PRSNet-2 and seven baseline methods across eight diseases. (B) The AUPRC scores of PRSNet-2 and baseline methods for different diseases. The bar plot represents the mean score, and the error bar indicates the standard error. (C) AUPRC and AUROC scores of PRSNet-2 and baseline methods for Alzheimer’s disease prediction on the independent target cohort, ADNI. (D) The cumulative incidence rates of eight diseases for high-risk (top 5%), median-risk (middle 5%), and low-risk (bottom 5%) individuals stratified based on PRSNet-2 prediction. The training, validation, and testing procedures were repeated five times with different random seeds for each model and each disease. AUPRC, the area under the precision-recall curve; AUROC, the area under the receiver operating characteristic curve; ADNI, AD Neuroimaging Initiative.

We next used the cumulative incidence rate curve to illustrate disease occurrence over a lifetime for individuals stratified into high-risk, median-risk, and low-risk groups based on PRSNet-2’s prediction. High-risk individuals were defined as those in the top 5% of PRSs, median-risk individuals as those in the middle 5%, and low-risk individuals as those in the bottom 5%. Individuals classified as high-risk exhibited a significantly elevated risk of disease throughout their lifetime, while low-risk individuals presented a consistently lower risk (Fig. 2D). These findings support the potential of PRSNet-2 as a powerful tool for disease risk stratification.

### PRSNet-2 enhances quantitative trait prediction

Next, we evaluated PRSNet-2 on two quantitative traits: height and body mass index (BMI), using the UKBB EUR samples for analysis. The coefficient of determination (*R*^2^) was utilized to assess predictive performance. PRSNet-2 demonstrated superior performance compared to all baseline methods for both traits, with relative improvements of 6.05% and 6.02% for height and BMI, respectively (Fig. 3A). To examine the robustness of these improvements, we repeated the analysis using a separate GWAS for the same trait. Again, PRSNet-2 surpassed all baselines, yielding relative *R*^2^ improvements of 7.42% for height and 5.93% for BMI. These results underscore the robustness and generalizability of PRSNet-2.

**Fig. 3:**
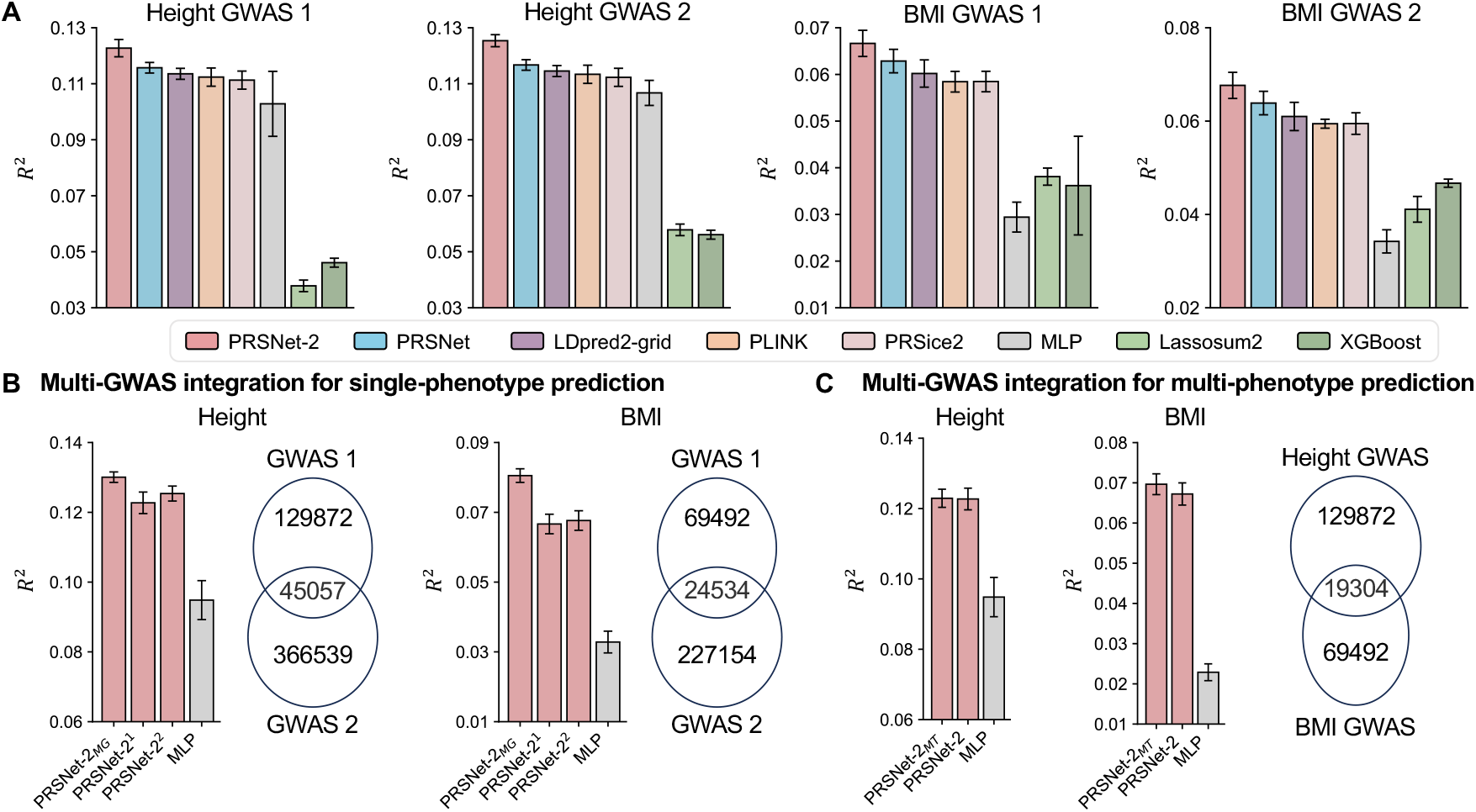
Predictive performance evaluation for quantitative traits. Performance was measured by the coefficient of determination (*R*^2^). (A) Performance of PRSNet-2 and baseline methods for height and BMI, where the results of two GWASs for each trait were presented. (B) Comparison of PRSNet-2 integrating two GWASs (PRSNet-2_MG_) versus using a single GWAS (PRSNet-2^1^ and PRSNet-2^2^). (C) Performance of PRSNet-2 integrating two GWASs for multitask learning of height and BMI (PRSNet-2_MT_) compared to the standard PRSNet-2 and MLP. The Venn plots illustrate the overlap of selected variants between the two GWASs. The bar plot and error bar denote the mean and the standard error, respectively. The training, validation, and testing procedure was conducted for five repeats with different random seeds for each model and each trait. BMI, body mass index.

We also assessed PRSNet-2’s capacity to integrate multiple GWASs for phenotype prediction. Our results show that PRSNet-2, when integrating two GWASs (denoted as PRSNet-2_MG_), outperformed the version based on a single GWAS, achieving relative improvements of 3.75% for height and 19.03% for BMI (Fig. 3B). By combining two GWASs, we obtained over 400K and 200K SNPs for height and BMI prediction, respectively. While traditional MLP models struggled to effectively utilize these large numbers of SNPs, PRSNet-2 managed to handle high-dimensional data without overfitting, resulting in superior performance (Fig. 3B). This underscores PRSNet-2’s ability to integrate diverse genetic signals, model high-dimensional features, and mitigate overfitting.

Finally, by integrating the GWASs for height and BMI, PRSNet-2 enabled multitask learning for both traits (denoted as PRSNet-2_MT_), exhibiting enhanced performance compared to the single-task version (Fig. 3C). Collectively, these findings demonstrate the generalizability and scalability of PRSNet-2 for diverse application scenarios.

### PRSNet-2 identifies disease-relevant genes

Unlike traditional PRS methods, the inherent interpretability of PRSNet-2 facilitates the identification of genes associated with diseases. To demonstrate this, we first identified genes with significant differences in their PRSNet-2 attention weights between RA disease and control groups. This analysis revealed 84 genes in total (adjusted *P* -value *<* 0.05 and |effect size| *>* 15% of reference effect sizes) (Fig. 4A).

**Fig. 4:**
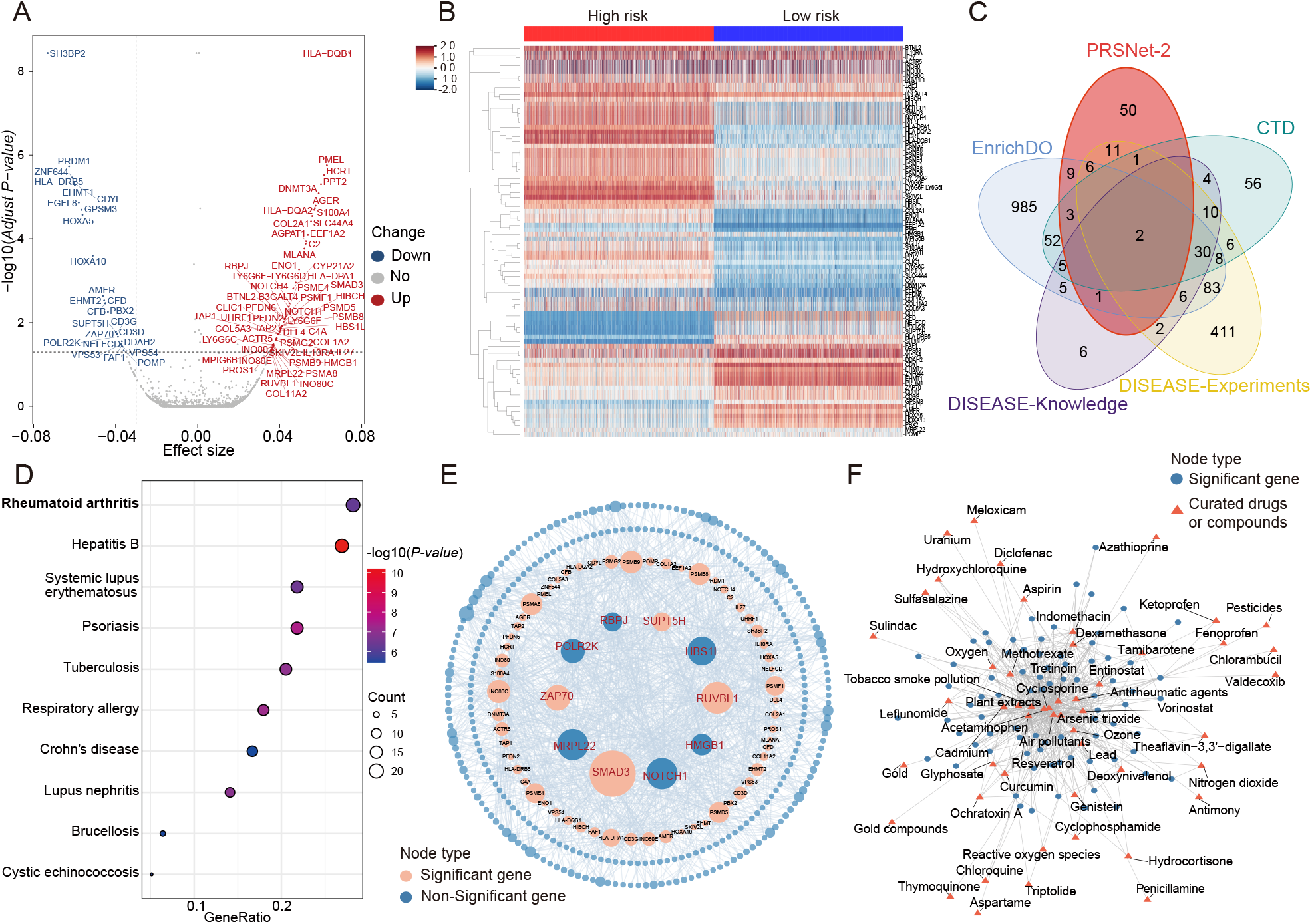
PRSNet-2 identifies disease-relevant genes for rheumatoid arthritis (RA). (A) Volcano plot of disease-relevant genes identified by PRSNet-2. The Mann-Whitney U test was conducted to identify significant RA-related genes. *P*-values were adjusted using the Benjamini-Hochberg (BH) procedure, and effect sizes were calculated using the Vargha-Delaney A statistic, which quantifies the effect associated with the U statistic. Red dots represent genes with significantly higher weights (adjusted *P*-value *<* 0.05) in patients, while blue dots represent genes with lower weights. Gray dots denote non-significant genes or genes within the bottom 15% of effect sizes. (B) Heatmap of attention weights for RA-related genes prioritized by PRSNet-2, comparing high-risk (top 5% of PRSNet-2 predictions) and low-risk (bottom 5% of PRSNet-2 predictions) individuals. The color gradient from blue to red indicates the range of gene weights from low to high. (C) Venn diagram illustrating the intersection between genes identified by PRSNet-2 and RA-associated genes collected from the comparative toxicogenomics database (CTD), DISEASE dataset, and EnrichDO dataset. (D) Results of weighted disease ontology analysis conducted using EnrichDO, based on genes identified by PRSNet-2. (E) Hub gene detection results based on the GGI network. A hub gene is defined as a gene with one of the top ten effect scores, where the effect score is calculated as |effect size| ×degree×betweenness. (F) Drug-gene interaction network (derived from the CTD database), which is constructed based on verified drug interactions for genes identified by PRSNet-2.

Next, we stratified individuals based on their PRSNet-2 predicted risk scores into high-risk (top 5%) and low-risk (bottom 5%) groups. Clustering analysis highlighted distinct differences in gene attention weights between these two groups (Fig. 4B). Notably, genes such as *DLL4, NOTCH1, SMAD3, NOTCH4*, and *RBPJ* exhibited higher attention weights in the high-risk group, suggesting their important role in impacting RA risk. These findings are indeed consistent with previous studies that have linked these genes to RA pathogenesis^[29,30,31,32,33]^.

We then cross-referenced the identified genes with those reported in three established gene-disease databases: the Comparative Toxicogenomics Database (CTD)^[34]^, DISEASE^[35]^, and EnrichDO^[36]^, which provide validated gene-disease associations, experimental support, and text-mining-derived gene-disease relationships. Among the 84 genes identified by PRSNet-2, 40.4% were supported by the literature or experimental evidence linking them to RA (Fig. 4C).

To further evaluate the biological relevance of the prioritized genes, we performed disease ontology (DO) enrichment analysis using the EnrichDO package, employing its weighted enrichment analysis algorithm based on the genes identified by PRSNet-2. This analysis revealed a significant enrichment for RA, confirming the disease relevance of the genes (Fig. 4D). Additionally, KEGG pathway enrichment analysis prioritized Th1 and Th2 cell differentiation pathways (Supplementary Fig. 2). Previous studies in mouse models have demonstrated that an imbalance in these pathways is crucial for the development of RA^[37]^, underscoring the model’s ability to identify genes central to RA’s pathophysiology.

We also conducted hub gene detection analysis using the GGI network. The top 10 genes with the highest effect scores were identified as hub genes, including *SMAD3, HMGB1, HBS1L, RUVBL1, SUPT5H, MRPL22, ZAP70, NOTCH1, RBPJ*, and *POLR2K*. Seven of these hub genes, such as *SMAD3* ^[31]^, *HMGB1* ^[38]^, *HBS1L* ^[39]^, *RUVBL1* ^[40]^, *ZAP70* ^[41]^, and *NOTCH1* ^[30]^, have previously been implicated in RA pathogenesis (Fig. 4E). Moreover, *RBPJ* has been identified as a known RA marker gene^[33]^.

Finally, we explored drug-gene interactions for the RA-related genes identified by PRSNet-2. Using data from the CTD database, we found that 83 out of the 84 genes interact with 932 compounds. We then queried these compounds for their association with RA. Thirteen compounds were found to be associated with RA or implicated in its etiology, while 38 compounds were identified as having known or potential therapeutic effects on RA (Fig. 4F). These results not only validate our identified genes but also suggest potential therapeutic targets, emphasizing the utility of PRSNet-2 in both identifying disease-related genes and uncovering potential drug candidates.

### Ablation studies

To examine the contribution of different components in PRSNet-2, we conducted extensive ablation studies. We first compared multiple variations of PRSNet-2, including versions without SG-Dropout, without SG-L1, and without both modules. PRSNet-2 consistently outperformed all these variants in Alzheimer’s disease prediction (Fig. 5A), with relative improvements of 4.36%, 12.81%, and 17.08% in AUROC, respectively. These results underscore the critical role of the significance-guided regularization strategies in enhancing prediction performance. Next, we inspected the effects of varying the number of kernels and the SG-L1 weight on the performance (Fig. 5B). Initially, increasing the number of kernels led to performance improvement; however, beyond a certain point, adding more kernels caused a decline in performance. The relatively low similarity between kernels (Fig. 5C) suggests that different kernels capture distinct genetic information, thereby enriching the gene-level representations. Similarly, increasing the SG-L1 weight initially boosted performance, but excessively large weights resulted in a drop in performance. Lastly, we assessed the impact of different *P* -value thresholds for variant selection in PRSNet-2. Overall, PRSNet-2 demonstrated robustness to variations in the *P* -value cutoff (Fig. 5D). For Alzheimer’s disease prediction, as the *P* -value threshold increased, the predictive performance of PRSNet-2 showed a positive trend, while both PLINK and MLP models exhibited a marked decline in performance. For BMI prediction, PRSNet-2, PLINK, and MLP all showed an increasing trend as the *P* -value threshold rose. However, when the *P* -value threshold was particularly large (e.g., *>* 0.1), PRSNet-2 outperformed the others, demonstrating superior prediction accuracy and a stronger ability to combat overfitting. This further suggests that PRSNet-2 is better at leveraging a broader range of genetic information while effectively avoiding overfitting compared to these traditional models.

**Fig. 5:**
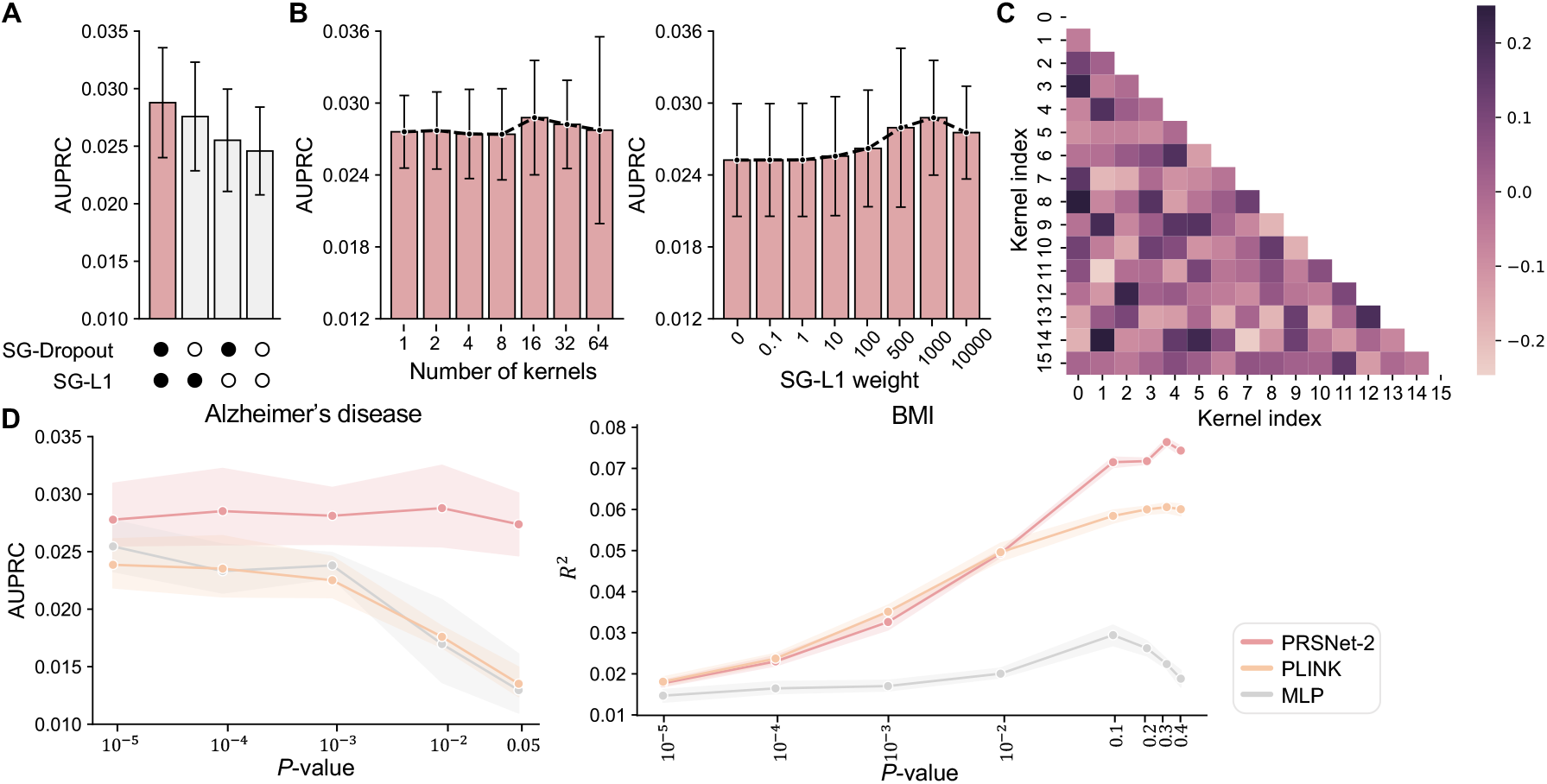
The ablation results for PRSNet-2. (A) Performance comparison for Alzheimer’s disease between PRSNet-2 and its variants, including models without SG-Dropout, without SG-L1, and without both SG-Dropout and SG-L1. The filled circle and hollow circle represent models with and without the corresponding modules, respectively. (B) Performance of PRSNet-2 for Alzheimer’s disease with varying numbers of kernels and different SG-L1 weight values. (C) Heatmap of cosine similarity between kernel weights for the PRSNet-2 model applied to Alzheimer’s disease. (D) Performance of PRSNet-2 for Alzheimer’s disease (left) and BMI (right) with different *P* -value thresholds used for SNP selection. The bar plot and error bar denote the mean and the standard error, respectively.

## Discussion

In this study, we introduce PRSNet-2, an interpretable, end-to-end deep learning framework specifically engineered for the prediction of complex phenotypes directly from high-dimensional genotypic data. PRSNet-2 employs a novel hierarchical GNN architecture to address the challenges inherent in high-dimensional genetic data. The key architectural upgrade in PRSNet-2 is the introduction of a multi-kernel aggregator. Unlike the original PRSNet, which utilized conventional C+T methods to calculate gene-level PRS, PRSNet-2 leverages the multi-kernel aggregator to learn a more flexible and comprehensive gene-level representation from the vast genotypic space. This shift enables PRSNet-2 to fully capture the complex interactions among SNPs, resulting in a more informative genetic embedding. To manage the massive scale and complexity of genetic data — especially its high dimensionality, polygenicity, and sparsity inherent in complex phenotype genetics — we developed significance-guided model regularization strategies including dropout and *L*_1_ penalty. These strategies proved essential to enhance PRSNet-2’s capacity to process approximately half a million genetic variants while successfully mitigating the issue of overfitting.

Extensive empirical evaluations across a wide range of complex traits and diseases demonstrated that PRSNet-2 substantially outperformed various established baseline methods, including the previous PRSNet. Furthermore, PRSNet-2 exhibits high versatility and extendability. It was shown to be easily adaptable to integrate multiple GWAS datasets into a single, unified model. This capability significantly enhanced predictive performance in both single-phenotype and multi-phenotype prediction tasks. The inherent interpretability of PRSNet-2 extends its utility beyond prediction. Specifically, in the context of rheumatoid arthritis, the framework successfully identified disease-relevant genes, functional gene modules, and potential therapeutic targets underlying the complex disease. The future work includes the integration of non-genetic features, such as environmental factors, lifestyle, and other omics data, into the PRSNet-2 framework to further enhance phenotype prediction. In summary, PRSNet-2 offers a powerful, versatile tool for both genetic risk stratification and biological insight discovery.

## Data availability

All genomic and phenotypic data used in this study are available from UKBB upon application (https://www.ukbiobank.ac.uk). The independent target cohort for AD are acessible through ADNI (https://adni.loni.usc.edu/data-samples/adni-data/).

## Software availability

The source code and the tutorial of PRSNet-2 are available at GitHub (https://github.com/lihan97/PRSNet-2).

## Acknowledgments

This work was supported by the National Key RD Program of China (2024YFC3407800 to H.L. and X.X.), Natural Science Foundation of Tianjin(25JCQNJC01180 to H.L.), the Sponsored by Beijing Nova Program, the National Natural Science Foundation of China (82573343 to X.X.), and the Strategic Priority Research Program of the Chinese Academy of Sciences (XDA0460203 to X.X.). This work was supported by the National Key R&D Program of China (2024YFC3407800 to H.L. and X.X.).

## Author contributions

H.L. and S.Z. conceived the concept and designed the study. H.L. developed PRSNet-2 and performed data analysis. H.L., Z.S., H.F., Y.D., S.G., P.G., J.C.-K., Y.M., X.X. and S.Z. are responsible for data interpretation. S.Z., X.X., Y.M and H.L. supervised the project. H.L. and S.Z. prepared the manuscript with assistance from all other authors.

